# Hippocampal trauma memory processing conveying susceptibility to traumatic stress

**DOI:** 10.1101/2024.01.08.574632

**Authors:** Bart C.J. Dirven, Lennart van Melis, Teya Daneva, Lieke Dillen, Judith R. Homberg, Tamas Kozicz, Marloes J.A.G. Henckens

**Affiliations:** Department of Cognitive Neuroscience, Donders Institute for Brain, Cognition, and Behaviour, Radboud University Medical Centre, 6500 HB, Nijmegen, The Netherlands; Department of Medical Imaging, Anatomy, Donders Institute for Brain, Cognition, and Behaviour, Radboud University Medical Centre, 6500 HB, Nijmegen, The Netherlands; Center for Individualized Medicine, Department of Clinical Genomics, and Biochemical Genetics Laboratory, Mayo Clinic, Rochester, MN 55905, Minnesota, USA; University of Pecs Medical School, Department of Anatomy, Pecs, Hungary

**Keywords:** Stress, Fear Memory, Resilience, Hippocampus

## Abstract

While the majority of the population is ever exposed to a traumatic event during their lifetime, only a fraction develops posttraumatic stress disorder (PTSD). Disrupted trauma memory processing has been proposed as a core factor underlying PTSD symptomatology. We used transgenic Targeted- Recombination-in-Active-Populations (TRAP) mice to investigate potential alterations in trauma-related hippocampal memory engrams associated with the development of PTSD-like symptomatology. Mice were exposed to a stress-enhanced fear learning paradigm, in which prior exposure to a stressor affects the learning of a subsequent fearful event (contextual fear conditioning using foot shocks), during which neuronal activity was labeled. One week later, mice were behaviorally phenotyped to identify mice resilient and susceptible to developing PTSD-like symptomatology. Three weeks post-learning, mice were re-exposed to the conditioning context to induce remote fear memory recall, and associated hippocampal neuronal activity was assessed. While no differences in the size of the hippocampal neuronal ensemble activated during fear learning were observed between groups, susceptible mice displayed a smaller ensemble activated upon remote fear memory recall in the ventral CA1, higher regional hippocampal PV^+^ neuronal density and a relatively lower activity of PV^+^ interneurons upon recall. Investigation of potential epigenetic regulators of the engram revealed rather generic (rather than engram-specific) differences between groups, with susceptible mice displaying lower hippocampal histone deacetylase 2 expression, and higher methylation and hydroxymethylation levels. These finding implicate variation in epigenetic regulation within the hippocampus, as well as reduced regional hippocampal activity during remote fear memory recall in interindividual differences in susceptibility to traumatic stress.

## Introduction

Posttraumatic stress disorder (PTSD) is a debilitating disorder one can develop after exposure to a traumatic event. One of the hallmark features of PTSD is the re-experiencing of the trauma by flashbacks, spontaneous recollections, and recurrent nightmares of the trauma, which affect over 90% of patients (DSM-5, Green, 2003). Behavioral treatment strategies in which the trauma memory is targeted are among the most effective clinical treatments for PTSD (Watkins et al., 2018, Wilson et al., 2018), implicating disrupted trauma memory processing in PTSD. Interestingly, whereas the majority of the population is ever exposed to a traumatic event during their lifetime, only a small fraction of them develops PTSD (Kessler et al., 2005). We hypothesize that resilience may be characterized by adaptive trauma memory processing, which turns maladaptive in susceptible individuals. During trauma processing, the complex configuration of trauma-related information triggers the activity of neural ensembles that communicate through neuronal synapses, which are subsequently strengthened and stabilized through synaptic plasticity at the neuronal and circuit level (Lacagnina et al., 2019). These neural ensembles in which the memory is physically stored are referred to as the memory engram (Josselyn and Tonegawa, 2020, Maddox et al., 2019). The development of new genetic tools provides current, unprecedented opportunities to capture and study these engrams (Josselyn et al., 2015). Here, we make use of Targeted Recombination in Active Populations (TRAP) to investigate whether PTSD-like symptomatology is associated with an aberrant trauma-related hippocampal memory engram.

Decades of work have implicated the hippocampus as an important site for memory engrams (Josselyn et al. 2015), through its role in contextual memory processing (Brohawn et al., 2010, Shin et al., 2006), and its modulation by the amygdala in case of emotionally salient events (Akirav and Richter-Levin, 2002, Richter-Levin and Akirav, 2000, Tsoory et al., 2008). Neuroimaging studies have observed smaller hippocampal volume (Logue et al., 2018) and impaired function (Shin et al., 2006) in PTSD patients, while animal models for PTSD have shown increased hippocampal apoptosis (Li et al., 2010), reduced levels of brain-derived neurotrophic factor (Kozlovsky et al., 2007) and increased glucocorticoid receptor expression (Knox et al., 2012), implicating aberrant hippocampal function in PTSD pathophysiology. Furthermore, reduced hippocampal activity during exposure to trauma-related stimuli has been positively correlated with PTSD severity (Astur et al., 2006) and trauma-related memory distortions in PTSD-affected combat veterans (Hayes et al., 2011). Yet, it remains unclear how these rather generic hippocampal abnormalities relate to potential deviations in the trauma memory engram. Here, we investigated whether deviations in the hippocampal fear memory engram code vulnerability to the long-term consequences of trauma exposure in terms of PTSD-like symptomatology in mice, dissociating ventral from dorsal hippocampus (Fanselow and Dong, 2010, Matus-Amat et al., 2004, McHugh et al., 2004), as well as hippocampal subregions (i.e., dentate gyrus (DG), Cornu Ammonis areas 1 (CA1) and 3 (CA3)) (Daumas et al., 2005).

As potential modulators of the engram, we investigated parvalbumin positive (PV^+^) interneurons, which innervate large numbers of hippocampal pyramidal neurons and are spatially well-positioned to coordinate neuronal ensemble activity (Hu et al., 2014). Their activity has been shown required for the stabilization of hippocampal connectivity networks upon learning of a novel experience (Ognjanovski et al., 2017), and PV^+^ neurons have been shown vulnerable to the effects of prolonged stress (Czeh et al., 2015, Filipovic et al., 2013). Additionally, we investigated epigenetic regulation, which confers transcriptional memory of exposure to environmental stress conditions (Fabrizio et al., 2019, Tsankova et al., 2007), regulates memory formation (Levenson et al., 2004) and shapes long-term behavioral adaptations (S. Jiang et al., 2019, Siegmund et al., 2007, Uchida et al., 2011). Histone acetylation is most robustly associated with memory formation (Graff and Tsai, 2013) and the expression of particularly hippocampal histone deacetylase (HDAC) 2 is negatively related to memory performance and hippocampal plasticity (Guan et al., 2009, Peixoto and Abel, 2013). Prior reports have shown that chronic stress downregulates hippocampal HDAC2 levels, causing depressive-like symptomatology in mice (Lee et al., 2019). Yet, others have reported on a stress protective effect of HDAC2 reductions (Covington et al., 2009, Wang et al., 2017). Similarly, stress exposure changes DNA methylation state (Matosin et al., 2017), with both stress-induced increases (Hammels et al., 2015, Sales and Joca, 2018) and reductions (Rodrigues et al., 2015) in hippocampal DNA methylation being observed. Also 5- hydroxymethylcytosine (5hmC) levels, a stable epigenetic modification (Bachman et al., 2014) modulating gene transcription independently from 5mC (Lin et al., 2017), have been shown to be modulated by prior stress exposure (Li et al., 2015).

We here used a mouse model to test our hypothesis that alterations in trauma-related hippocampal engrams are associated with the development of PTSD-like symptomatology and investigated aforementioned key engram regulators potentially at the core of these alterations. The PTSD mouse model used is based on the phenomenon of stress-enhanced fear learning (SEFL (Rau et al., 2005, Rau and Fanselow, 2009)), with prior stress exposure affecting fear learning and memory. Mice were therefore first exposed to a stressor (severe, uncontrollable, unpredictable foot shocks), followed by contextual fear conditioning (mild foot shock) the next day. Critically, in this PTSD model, mice were behaviorally tested for PTSD-like symptoms to dissociate susceptible from resilient mice (Dirven et al., 2022a, Dirven et al., 2022b, Henckens et al., 2017, Lebow et al., 2012, Preston et al., 2020) and delineate distinct SEFL memory formation and recall in these subgroups, respectively. Engram neurons activated during the encoding of SEFL were identified by using the TRAP transgenic mouse model (Guenthner et al., 2013), whereas those supporting remote fear memory recall were identified by conditioning context re-exposure three weeks later by immunohistochemistry for the immediate early gene cFos. PV^+^ interneuron presence and activity, as well as HDAC2, 5mC and 5hmC expression levels in both engram and non-engram neurons (i.e., neurons activated neither during fear memory encoding nor recall) were assessed by immunohistochemistry as well.

## Materials & Methods

### Animals

Two founder mouse lines, ArcCreER^T2^ (B6.129(Cg)-*Arc^tm1.1(cre/ERT2)Luo^*/J) and conditional tdTomato (B6.Cg-*Gt(ROSA)26Sor^tm9(CAG-tdTomato)Hze^*/J, 007909), were purchased from The Jackson Laboratory and bred as described before (Guenthner et al., 2013) to generate heterozygote ArcCreER^T2^xROSA offspring, referred to as ArcTRAP. This genetic construct allows *Arc*-expressing (i.e., active) neurons to be labeled by the fluorescent protein tdTomato in a 36-hour time window after injection with the compound tamoxifen. ArcTRAP mice were prefered over the available FosTRAP mice based on their superior labeling sensitivity in the hippocampal CA3 and CA1, which are typically devoid of labeled cells in the FosTRAP mouse lines (Dirven et al., 2022b, Guenthner et al., 2013). Because the PTSD

model (Dirven et al., 2022a, Dirven et al., 2022b, Guenthner et al., 2013, Henckens et al., 2017, Lebow et al., 2012) has only been validated in males, experiments were restricted to male mice. Mice were group housed (3-4 mice per cage) in individually ventilated cages on a reverse 12 h light/dark cycle (09:00 - 21:00 h) at the Central Animal Facility of the Radboud University Nijmegen, The Netherlands, according to institutional guidelines. Food and water were provided *ad libitum*. Unless otherwise stated, behavioral testing was performed during the animal’s active phase (i.e., the dark) between 13.00 - 18.00 1. h. The experimental protocols were in line with international guidelines, the Care and Use of Mammals in Neuroscience and Behavioral Research (National Research Council 2003), the principles of laboratory animal care, as well as the Dutch law concerning animal welfare and approved by the Central Committee for Animal Experiments, Den Haag, The Netherlands.

### General procedure

44 ArcTRAP mice were injected with tamoxifen to induce fluorescent labeling of all *Arc*-expressing neurons and subsequently exposed to a PTSD mouse model as described before (Henckens et al., 2017, Lebow et al., 2012) (Figure 1A). The model is based on stress-enhanced fear learning (SEFL), which builds on the clinical observation that prior stress exposure precipitates PTSD (Boasso et al., 2015, Delahanty et al., 2003). Interestingly, SEFL has been found to robustly affect the learning of future aversive events, arguably in a primarily adaptive manner, creating stronger fear memories with increased resistance to extinction in general (Maren and Holmes, 2016, Rau et al., 2005, Rau and Fanselow, 2009, Sillivan et al., 2017), Yet, in addition, it has been shown to induce persistent anxiety- and arousal-related behavioral symptoms (i.e., impaired risk assessment, increased marble burying, shortened latencies to startle, impaired pre-pulse inhibition and increased locomotor activity in the inactive (light) phase of the circadian cycle) in a specific subset of susceptible mice (Dirven et al., 2022a, Dirven et al., 2022b, Henckens et al., 2017, Lebow et al., 2012, Preston et al., 2020). The behavioral profile of the susceptible animals resembles observations in PTSD patients, while they also display the hypothalamic-pituitary-adrenal (HPA)-axis abnormalities as observed in subgroups of PTSD patients (i.e., reduced glucocorticoid peak levels upon challenge mediated by increased negative feedback (Yehuda, 2009)). Importantly, susceptible mice do not show a different behavioral phenotype prior to SEFL (Dirven et al., 2022a), suggesting differential, supposedly maladaptive, responding to the SEFL procedure. The behavioral and neuroendocrine consequences are not observed if mice are only exposed to the initial stressor (Lebow et al., 2012), emphasizing deviations in subsequent fear learning to be at the core of the development of symptomatology.

**Figure 1.**
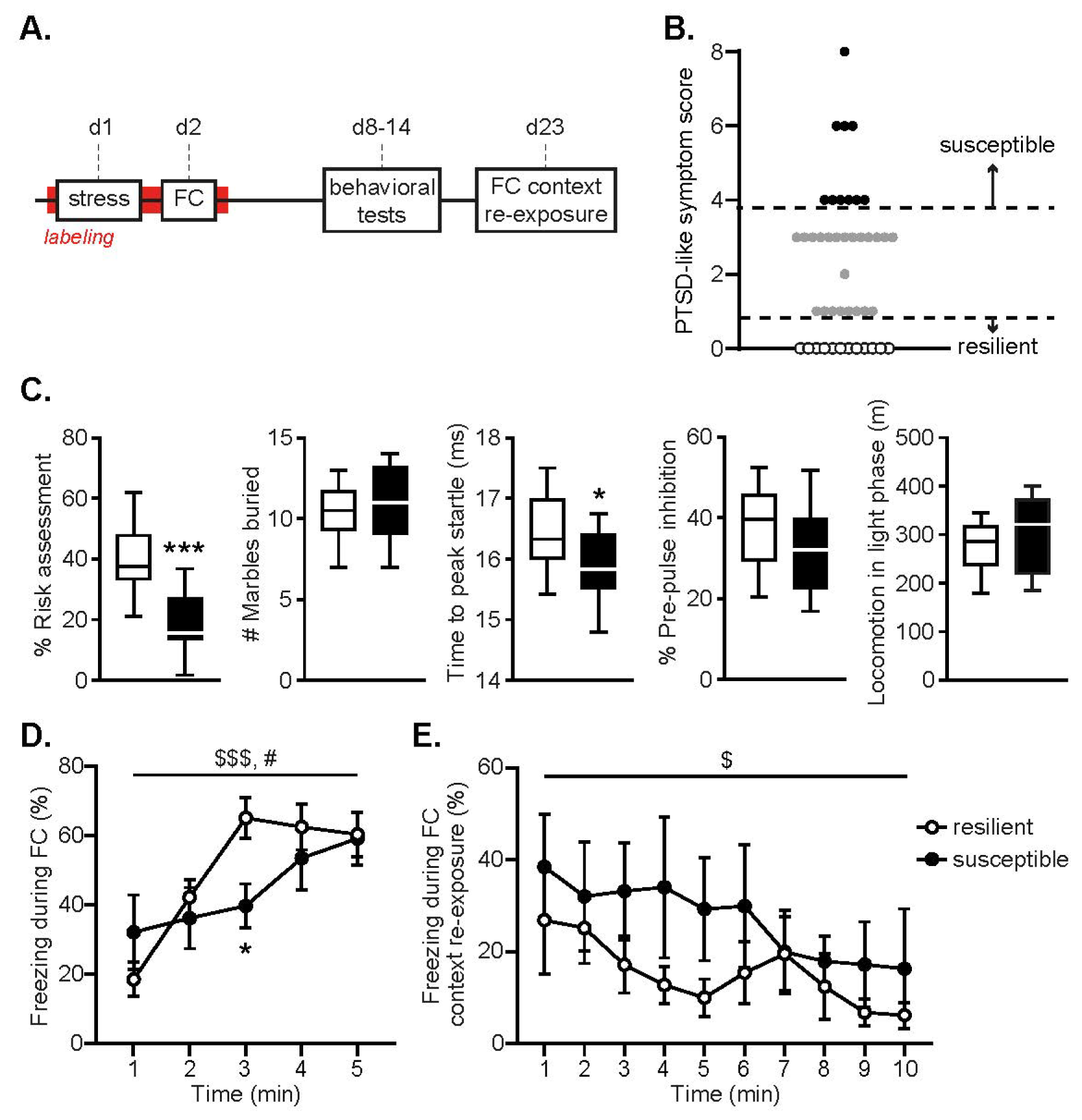
Experimental schedule and behavioral assessments. Mice were exposed to a stressor and then subjected to contextual fear conditioning (FC). PTSD-like symptomatology was assessed in a set of behavioral tests, mice were re-exposed to the conditioning context and then sacrificed (**A**). Susceptible mice were defined by PTSD-like symptom scores >=4 (necessitating extreme behavior in multiple tests), whereas resilient mice did not show any aberrant behavior (score = 0) (**B**). Susceptible mice displayed significantly reduced risk assessment behavior and shorter latencies to peak startle compared to resilient animals at the group level. No group differences were observed in marble burying, pre-pulse inhibition or locomotor activity during the light phase (**C**). Increases in freezing behavior during contextual FC differed across groups (shock administration at 1, 2, 3, 4 and 5 min) (**D**), whereas no differences in freezing levels were observed upon later FC context re-exposure (no shocks) (**E**). Data are presented as medians with interquartile ranges (**B**) and mean +/- standard error of the mean (**C**). $$$: p < .001, $: p < .05, main effect of time; #: p < .05, group x time interaction; ***: p < .001, *: p < .05 effect of group.

To induce a PTSD-like phenotype in susceptible mice, all mice were exposed to an initial stressor, followed by fear learning (contextual fear conditioning) the next day. After a week, mice were subjected to a subset of behavioral tests over the course of two weeks to assess PTSD-like symptomatology. One week after the final behavioral test, mice were re-exposed to the conditioning context for 10 minutes to induce fear memory recall and sacrificed by perfusion-fixation 90 minutes later.

### Tamoxifen

Tamoxifen was dissolved in a 10% ethanol / corn oil solution at a concentration of 10 mg/mL by overnight sonication and stored at -20°C until further use. Solutions were heated to body temperature and intraperitoneally injected at a dosage of 150 mg/kg to induce activity-dependent neuronal labeling. Mice were injected with tamoxifen on the morning of day 1 - seven hours before the stressor - to induce SEFL-dependent active neuronal labeling. Fear learning was conducted 21 hours post-stressor. This allowed both the stressor and fear learning to fall within the 36-hour labeling window, capturing neuronal activation during both events. We opted for this approach as it is currently unknown whether the interindividual differences in SEFL and its long-term consequences originate from differential responding to the first stressor or from later fear learning. We hypothesized the latter, as the behavioral consequences of this PTSD-model are not observed to a similar degree if mice are only exposed to the initial stressor (Lebow et al., 2012), emphasizing aberrant fear learning to be at the core of the development of symptomatology. However, others have shown that PTSD-like memories can also be induced by stress exposure post-learning (Al Abed et al., 2020, Kaouane et al., 2012) (though applied immediately post-learning rahter than delayed), leaving this issue unresolved. Moreover, injection of 4- hydroxytamoxifen instead of tamoxifen to ensure a more specific (∼12-hour) labeling window, comes at the downsite of inducing instant labeling of activated neurons, capturing the neurons processing the injection stress as well, increasing noise. Therefore, we opted to inject tamixfen instead, and labeling of neuronal activity that was non-SEFL-related was minimized by keeping the mice undisturbed in their home cage during the rest of the labeling period.

### PTSD protocol

Seven hours after the tamoxifen injection, mice were individually placed in *Context A* boxes, in which they received 14 1 second 1.0 mA shocks (i.e., the stressor) over 85 minutes in variable intervals. Mice were first moved to the dark experimental room in groups of two to three animals in dark carton boxes before being placed in the fear-conditioning boxes, which were connected to a shock generator (Campden Instruments). *Context A* consisted of a black, triangular shaped Plexiglas box with a steel grid and metal tray. The boxes were sprayed with 1% acetic acid, not illuminated, and 70 dB background noise was presented. Boxes were equipped with infrared beams at both ends, and beam break data was used to analyze gross locomotor activity during stress exposure.

On the second day, 28 hours after the tamoxifen injection, mice were individually placed in *Context B* boxes, in which they received 5 1 second shocks of 0.7 mA over a period of five minutes (i.e., the fear learning), presented over fixed intervals. For this trigger session, mice were moved to the 70 lux illuminated experimental room in see-through cages in groups of two to three animals. The *Context B* boxes contained curved white walls and a steel grid with a white tray underneath. The boxes were furthermore cleaned with 70% ethanol and during the session the house lights in the boxes were turned on. No background noise was presented. As such, all sensory features (olfactory, auditory, and visual) differed between contexts A and B, sharing the sensory input of the grid floor as only commonality.

Mice were allowed to recover for a week, after which their behavioral response to trauma was assessed by testing for PTSD-related behavior: impaired risk assessment (dark-light transfer test), increased threat-induced anxiety (marble burying), hypervigilance (acoustic startle), impaired sensorimotor gaiting (pre-pulse inhibition), and disturbed circadian rhythm (locomotor activity during the light phase) (Lebow et al., 2012).

### Behavioral testing

*Dark-light transfer test*. On day 8 of the protocol, tested in the dark-light transfer test. The test was executed in a box that was divided into a dark compartment (DC, 29 x 14 cm) and brightly illuminated (ca. 1100 lux) compartment (LC, 29 x 29 cm), connected by a retractable door. The mice were individually placed in the DC, and the door was opened to initiate a 5-minute test session. Movement of the mice was recorded and scored automatically with Ethovision XT (Noldus IT). An additional area of 6 x 3 cm surrounding the opening of the LC was programmed into the software tracking measurements. Time spent in the LC as well as time spent in this ‘risk assessment’ zone were measured. Percentage risk assessment was calculated as the amount of time spent in the risk assessment zone as a percentage of total time spent in the LC.

*Marble burying*. On day 10, mice were individually placed in a 10 lux illuminated black open box (30 x 28 cm), containing a 5 cm deep layer of corn cobs, on top of which 20 marbles were centrally arranged in a 4 x 5 grid formation. Each mouse was placed in the corner of the box to initiate the task. Mice were videotaped for 25 minutes. Videos were scored by assessing the number of buried marbles after 25 minutes.

*Startle response and pre-pulse inhibition*. On day 12, mice were moved to the experimental room in their home cage and individually placed in small, see-through Plexiglas constrainers mounted on a vibration-sensitive platform inside a ventilated cabinet that contained two high-frequency loudspeakers (SR-LAB, San Diego Instruments). Movements of the mice were measured with a sensor inside of the platform. The pre-pulse inhibition test (PPI) started with an acclimatization period of 5 minutes, in which a background noise of 70 dB was presented, which was maintained throughout the entire 30-minute session. Thirty-two startle cues of 120 dB, 40 ms in duration and with a random varying ITI (12-30 s), were presented with another 36 startle cues preceded by a 20 ms pre-pulse of either 75 dB, 80 dB or 85 dB. Sessions were scored by assessing the latency to peak startle amplitude of the 12 middle startle trials, and the pre-pulse inhibition, i.e., the percentage of startle inhibition response to the different pre-pulse stimuli [1 - (mean pre-pulse startle response / mean startle response without pre-pulse) x 100].

*Homecage locomotion*. Immediately after the pre-pulse inhibition test, mice were individually housed in Phenotyper cages (45 x 45 cm, Noldus) for 72 hours while their locomotion was being recorded by an infrared-based automated system (EthoVision XT, Noldus). The first 24 hours were considered habituation time and data were discarded. Total locomotion time during the subsequent two light phases (21:00 - 09:00 h) was assessed.

### Behavioral categorization

In order to categorize mice as either susceptible or resilient, one compound measure was generated based on the five behavioral outcome scores. Mouse behavior on each of the tests was sorted, and the 20% of mice that had the lowest values were attributed 3 points for percentage risk assessment, 3 points for latency to peak startle amplitude, and 2 points for percentage PPI. Similarly, the 20% of mice showing the highest values were attributed 1 point for light locomotor activity and marble burying (Henckens et al., 2017). Points for each test were determined by factor analysis in which tests were clustered in three separate groups: (1) latency to peak startle amplitude and percentage risk assessment, (2) percentage PPI, and (3) marble burying and total light activity (Lebow et al., 2012) (Table 1). Ties in the marble burying test were resolved by also assessing the number of marbles buried after 15 minutes, and assigning points to the mice that buried most marbles then. The points per animal were tallied to generate and overall PTSD-like symptom score. Mice that had a total of four or more points (necessitating extreme behavior in multiple tests) were termed susceptible. Only mice that had zero points (indicating no abnormal behavior within any of the tests) were termed resilient.

**Table 1.** Behavioral classification of resilient and susceptible mice. The 20% of mice that displayed the strongest phenotype on each test was assigned points, and points were tallied to generate a total PTSD-like symptom score.

| Behavioral measure | Points |
| --- | --- |
| *Low risk assessment* | 3 |
| *High marble burying* | 1 |
| *Short latency to peak startle* | 3 |
| *Low pre-pulse inhibition* | 2 |
| *High locomotor activity during the light phase* | 1 |
| **Total PTSD-like symptom score** | Resilient: 0<br>Susceptible: $\geq 4$ |

### Re-exposure and sacrifice

On the final day of the experiment, day 23, mice were re-exposed to the *Context B* box (i.e., the fear conditioning context) for 10 minutes to induce fear memory recall, following the exact same procedures as during the fear conditioning session. However, no shocks were administered during this context re-exposure session. Mice were sacrificed 90 min post re-exposure under anesthesia (5% isoflurane inhalation followed by intraperitoneal injection of 200 μL pentobarbital) by perfusion with phosphate buffered saline (PBS) followed by 4% paraformaldehyde solution (PFA). The brains were surgically removed and post-fixed for 24 hours in 4% PFA, after which they were transferred to 0.1 M PBS with 0.01% sodium azide and stored at 4°C.

### Freezing behavior

Mice were videotaped during fear conditioning (day 2) and the re-exposure to the conditioning context (day 23) to assess fear memory encoding and remote recall. Freezing behavior was manually scored by an observer blinded to the experimental condition using The Observer XT12 software (Noldus). Consistent with previous literature, mice were considered to freeze when they were immobile for more than two consecutive seconds (Patel et al., 2014, Shoji et al., 2014).

### Immunofluorescence

Right hemispheres of susceptible (n = 10) and resilient (n = 12) animals were sliced at 30 µm thickness using a freezing sliding microtome (Microm HM440E, GMI Inc., Ramsey, MN, USA) and stored in PBS with 0.01% sodium azide. Floating sections were used for immunohistochemistry of the hippocampus. For each animal, 4-6 sections were collected between anterior-posterior coordinates -1.46 mm and -1.94 mm relative to Bregma for the dorsal hippocampus, and between -2.92 mm and -3.52 mm relative to Bregma for the ventral hippocampus. tdTomato, as a proxy for the immediate early gene Arc, was used to measure neuronal activity during SEFL, while cFos immunofluorescence was assessed to measure recall-related activity. We used cFos, rather than Arc, because Arc labeling is primarily dendritic in some hippocampal subregions (Denny et al., 2014) complicating quantification of activated neurons, and both cFos and Arc expression have earlier been found to strongly overlap in neurons (Mahringer et al., 2019, Nakagami et al., 2013) - and specifically in the hippocampus (Jiang and VanDongen, 2021, Stone et al., 2011) - in response to a challenge.

*Immunolabeling of cFos and parvalbumin (PV) or histone deacetylase (HDAC) 2*. Sections were washed three times in 1x PBS and blocked in PBS-BT (1x PBS with 0.3% Triton X-100 and 1% bovine serum albumin) for 30 minutes at room temperature (RT). Incubation of the primary antibodies was performed overnight (guinea pig anti-cFos, 1:750, 226004, Synaptic Systems; rabbit anti-PV, 1:1000, ab11427, ITK; or rabbit anti-HDAC2, 3 µg/µL, AB_2533908, Thermo Fisher) in PBS-BT for 18 hours at RT. Then, sections were washed three times in 1x PBS, and incubated with the secondary antibodies (Alexa 647-conjugated donkey anti-guinea pig, 1:200, AP193SA6, Merck Chemicals; Alexa 488-conjugated donkey anti-rabbit, 1:200, A-21206, Thermo Fisher) in PBS-BT for 3 hours at RT. Lastly, slices were washed three times in 1x PBS, mounted on gelatin-coated slides using FluorSave^TM^ reagent (345789, Merck Chemicals) and cover slipped. The slices were stored at -20°C until image acquisition and cell counting.

*Immunolabeling of cFos, 5-methylcytosine (5mC) and 5-methylhydroxycytosine (5hmC)*. Sections were washed three times in 1x PBS and permeabilized in 1x PBS with 0.1% Triton X-100 for 5 minutes at RT. Then, slices were incubated in 1 M HCl for 2 hours, washed three times in 1x PBS and blocked in PBS-NT (1x PBS with 0.3% Triton X-100 and 8% normal goat serum) for 50 minutes, all at RT. Because this process bleaches endogenous fluorescence - here the tdTomato fluorescent signal - these slices had to be immunolabeled for red fluorescent protein (RFP) in addition to the other markers. After again washing the slices three times in 1x PBS, incubation of the primary antibodies was performed overnight (guinea pig anti-cFos, 1:750, 226004, Synaptic Systems; rat anti-RFP, 1:1000, 5f8, Chromotek; mouse anti-5mC, 1:500, GWB-BD5190, GenWay Biotech; rabbit anti-5hmC, 1:1000, AB_10013602, Active Motif) in PBS-NT for 18 hours at 4°C. Then, sections were washed three times in 1x PBS, and incubated with the secondary antibodies (Alexa 647-conjugated donkey anti-guinea pig, 1:200, AP193SA6, Merck Chemicals; Alexa 555-conjugated donkey anti-rat, 1:200, ab150154, Abcam; Alexa 488-conjugated goat anti-mouse, 1:200, A11001, Thermo Fisher; Alexa 405-conjugated anti-rabbit, ab175651, Abcam) in PBS-NT for 2 hours at RT. Lastly, slices were washed three times in 1x PBS, mounted on gelatin-coated slides using FluorSave^TM^ reagent (345789, Merck Chemicals) and cover slipped. The slices were stored at -20°C until image acquisition and cell counting.

### Image acquisition and cell counting

Images of the tdTomato/cFos/PV and tdTomato/cFos/HDAC2 signals were captured through a light microscope (Axio Imager 2, Zeiss) using a 10x (for tdTomato/cFos/PV) or 40x (for tdTomato/cFos/HDAC2) objective lens and a LED module (Colibri 2, Zeiss). Images of the tdTomato/cFos/5mC/5hmC staining were captured through a confocal microscope (LSM900, Zeiss) using a 40x objective lens. For the tdTomato/cFos/PV signal, as well as the tdTomato/cFos/HDAC2 signal, whole hippocampi were photographed. For the tdTomato/cFos/5mC/5hmC staining, the entire DG was photographed, while for the CA1 and CA3 regions three representative photos each were taken, with locations being consistent across slices and animals (Figure S1). Separate photos were stitched and cFos^+^, tdTomato^+^ and PV^+^ cells were manually counted per region in Fiji software (Schindelin et al., 2012) by an experimenter blinded to the experimental group. Hippocampal surface areas in each slice were assessed and corrected for to obtain standardized measures of cell density. Normalized cell counts were averaged per hippocampal subregion per animal and subjected to statistical testing. Note that the CA2 and CA1 regions were segmented together. This combined region will henceforth be referred to as ‘CA1’.

### Fluorescent signal intensity analysis

Expression levels of HDAC2, 5mC and 5hmC per cell were assessed by measuring signal intensity, and four cell clusters were identified by masks per hippocampal subregion per slice (Schindelin et al., 2012): 1) all tdTomato^+^cFos^-^ cells, 2) all cFos^+^tdTomato^-^ cells, 3) all tdTomato^+^cFos^+^ cells, and 4) all tdTomato^-^ cFos^-^ DAPI^+^ cells. Furthermore, a mask was generated for the background signal, which was obtained by inverting the DAPI^+^ mask. Within mask 1-4, the mean signal intensity of HDAC2, 5mC and 5hmC was assessed. Here, masks 1-3 define the fear memory engram cells, while mask 4 defines the non-engram cells. In the background mask, the mean of the signal intensity of HDAC2, 5mC and 5hmC was assessed to exclude potential inter-slice differences in background intensity. Analyses revealed that background HDAC2 signal was very consistent across slices and did not differ across the hippocampal axis (*F*(1,46.241) = 1.113, *p* = .297), hippocampal subregions (*F*(2,42.178) = .698, *p* = .503) or groups (group main effect; *F*(1,19.324) = .007, *p* = .936, all group interactions; *p*’s > .508). Background 5mC levels did depend on the hippocampus subregion (*F*(2,40.945) = 9.363, *p* < .001), but not axis (*F*(1,58.531) = .022, *p* = .884) or group (group main effect; *F*(1,9.793) = 1.994, *p* = .189, all group interactions; *p*’s > .068). A similar pattern was observed for background 5hmC levels that depended on hippocampal subregion (*F*(2,31.234) = 4.589, *p* = .018), but not axis (*F*(1,42.726) = 3.509, *p* = .068) or group (group main effect: *F*(1,14.919) = .496, *p* = .492, all group interactions: *p*’s > .832). Since no potentially confounding effects of background signals were detected, fluorescent signals were not background-corrected.

### Statistical Analyses

Data were analyzed using IBM SPSS Statistics 23. Normality was checked using the Shapiro-Wilk test. One resilient animal displayed deviant behavior (deviating more than two standard deviations from the groups’ mean) during the acoustic startle test, and was excluded from further analyses of the latency to peak startle and pre-pulse inhibition. For normally distributed data, statistical testing was performed by independent t-tests or one-way ANOVAs. Freezing behavior over time was analyzed by repeated measures ANOVA (with time as within-subjects factor, and group as between-subjects factor), whereas immunohistochemistry data was analyzed using linear mixed modelling implementing the restricted maximum likelihood estimation. In the latter, the factors axis (dorsal, ventral) and region (DG, CA3, CA1) were included as within-subjects variables, group as between-subjects variable and mouse as random intercept. For the epigenetic data, the factor engram type (non-engram (tdTomato^-^cFos^-^), encoding (tdTomato^+^), recall (cFos^+^), reactivated (tdTomato^+^cFos^+^) engram) was additionally included as within-subjects variable. For non-parametric data, the Mann-Whitney U test or Kruskal-Wallis test were used. Differences were considered statistically significant if *p* < 0.05.

## Results

### Behavioral differences between susceptible and resilient animals

To assess potential differences in hippocampal trauma-related engram activity associated with differential susceptibility to PTSD-like symptoms, a cohort of 44 ArcTRAP mice was exposed to the PTSD induction protocol. Following a week of recovery, mice were assessed for PTSD-like symptomatology to yield a group of susceptible (n = 10) and resilient animals (n = 12), which significantly differed on their overall PTSD-like symptom score (*U* = 120, *p* < .001) (Figure 1B). Symptomatology was rather heterogeneous across susceptible animals (Figure 1C), sharing some symptoms (percentage risk assessment (*t*(19) = 4.280, *p* < .001) and reaction time to peak startle (*t*(18) = 2.110, *p* = .025)), yet differing on others (marble burying (*t*(20) = .739, *p* = .234), percentage pre-pulse inhibition (*t*(17) = 1.210, *p* = .121) and locomotor activity in the light phase (*t*(14.633) = .864, *p* = .201). Thus, individual symptom profiles across susceptible mice differed.

Behavior during stress exposure was checked by assessing beam break data as proxy for locomotor activity. Susceptible and resilient mice did not differ in their overall locomotor activity during the stressor (*F*(1,14) = .041, *p* = .843), nor in its reduction over time (main effect of time: *F*(9.490, 132.857) = 25.682, *p* < .001, group x time interaction: *F*(9.490, 132.857) = 1.022, *p* = .427), indicating no gross behavioral differences during initial stress exposure. During the subsequent fear learning session, no overall group differences were observed in freezing rates (*F*(1,18) = .629, *p* = .438), yet the increase in freezing behavior over time (*F*(4,72) = 13.534, *p* < .001) significantly differed across groups (*F*(4,72) = 3.172, *p* = .019) (Figure 1D). Freezing levels tended to start lower in resilient mice, but also seemed to plateau sooner. *Post hoc* tests revealed only significant differences in the third minute of the fear learning session, when resilient mice displayed higher freezing levels than susceptible mice (*t*(18) = 2.870, *p* = .010). Freezing behavior upon re-exposure to the fearful context - to induce fear memory recall - was not different between resilient and susceptible animals (Figure 1E). Neither overall freezing levels (*F*(1,19) = 1.308, *p* = .267), nor the observed reduction in freezing over time (*F*(3.542,67.297) = 3.323, *p* = .019) differed between groups (group x time interaction: *F*(3.542,67.297) = .703, *p* = .576).

### Susceptible animals show a smaller activated neuronal ensemble within the vCA1 upon fear memory recall, but not during encoding

In the ArcTRAP mice, the neuronal ensemble active during SEFL, i.e., those neurons expressing the immediate early gene *Arc*, was permanently labelled by the reporter gene *tdTomato* (Figure 2ABC). No significant differences in the total number of activated hippocampal neurons during SEFL were observed between susceptible and resilient mice (*F*(1, 33.255) = .715, *p* = .404), nor was there any interaction effect between group and axis (*F*(1,40.938) = .880, *p* = .354), group and hippocampal subregion (*F*(2,30.033) = .295, *p* = .747) or group x axis x subregion interaction (*F*(2,30.033) = .255, *p* = .776), suggesting that hippocampal activity between groups was not different during initial memory formation. Neuronal activity associated with remote fear memory recall was measured by immunolabelling cFos^+^ neurons (Figure 2ABD); cells that were active during remote fear memory recall induced by re-exposure to the conditioning context. For the number of hippocampal neurons active upon recall, a significant group x hippocampal subregion interaction (*F*(2,20.586) = 5.055, *p* = .016) was found, and a trend towards a main effect of group (*F*(1, 4.034) = 4.100, *p* = .051). All other group interaction effects failed to reach significance (all *p*’s > .577). These effects were caused by lower neuronal activity during remote fear memory recall in the CA1 of susceptible vs. resilient animals (*F*(1,14.693) = 5.298, *p* = .036), most notably within the vCA1 (vCA1; *p* = .013, dCA1; *p* = .100).

**Figure 2.**
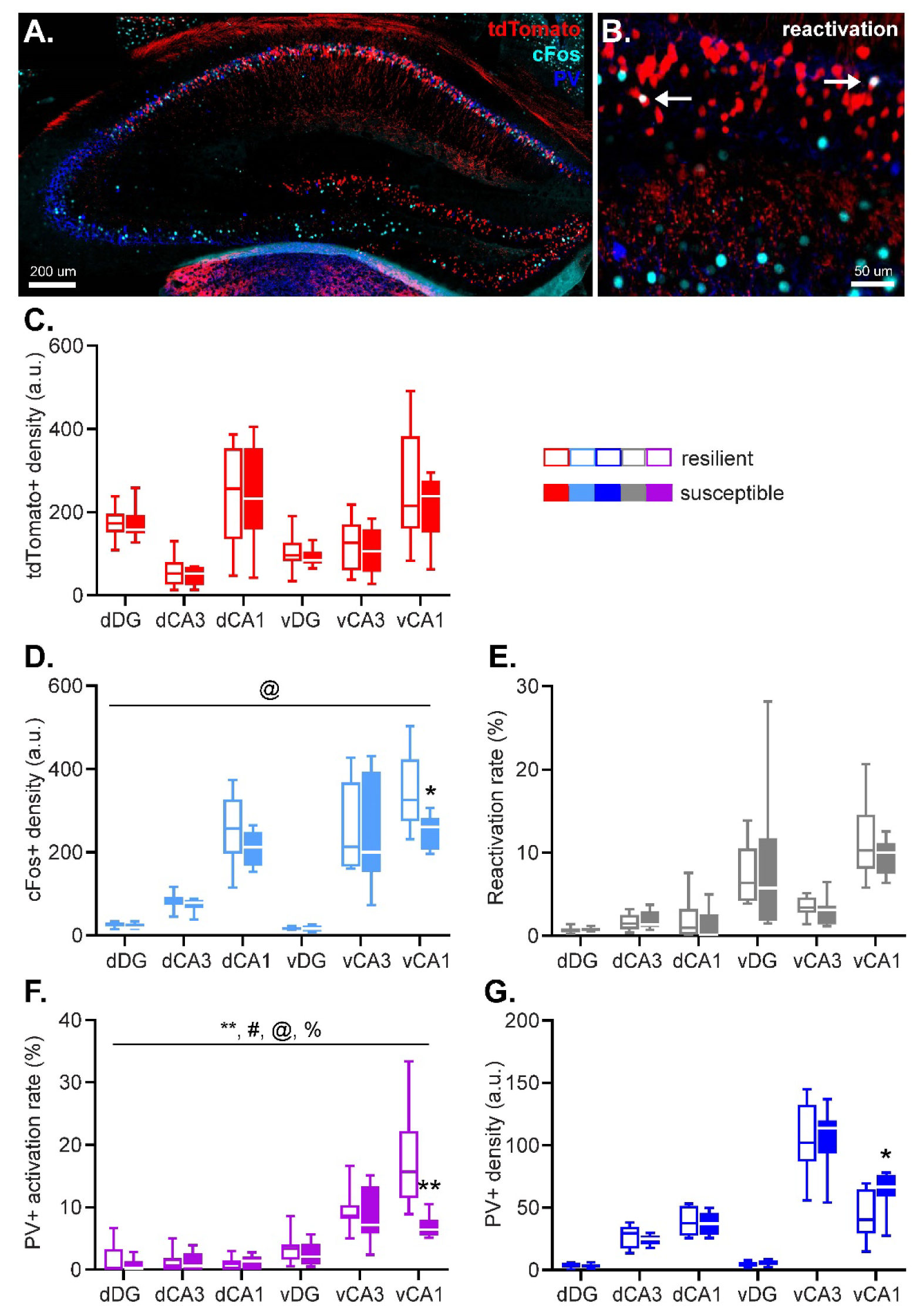
Hippocampal activity during fear memory encoding (marked by tdTomato expression), remote fear memory recall (marked by cFos expression), as well as parvalbumin (PV) interneuron density were assessed by immunohistochemistry. Arrows indicate tdTomato^+^cFos^+^ double-positive cells. (**AB**). No group differences were observed in the size of the engram recruited during stress-enhanced fear learning (**C**). However, susceptible animals displayed a smaller population of ventral hippocampal CA1 neurons active during remote fear memory recall (**D**), without any group differences in neuronal reactivation rate (**E**). Additionally, susceptible animals showed a decrease in activity of PV^+^ neurons in the ventral hippocampal CA1 specifically during remote memory recall (**F**), which was joined by an increase in local PV^+^ density (**G**). Data represent medians with interquartile ranges. **: p < .01, *: p < .05, main effect of group, @: p < .05, group x subregion interaction, %: p < .01, group x axis interaction, #: p < .05, group x axis x subregion interaction.

### Susceptible and resilient animals show no difference in hippocampal remote fear memory reactivation

To investigate which encoding-related (i.e., tdTomato^+^) cells eventually remained incorporated in the hippocampal memory engram for the fearful experience, overlap between the tdTomato^+^ and cFos^+^ neurons was assessed. These overlapping signals represent neurons that were active both during trauma encoding and recall, and therefore reflect the stable memory trace. Neuronal reactivation is expressed as the Reactivation Rate (RR), which is calculated by dividing the number of cFos^+^tdTomato^+^ overlapping neurons by the number of tdTomato^+^ neurons (Cowansage et al., 2014, Milczarek et al., 2018).

An average of 4.2% of hippocampal tdTomato^+^ neurons were reactivated during the trigger context re- exposure, with RRs in the different subregions ranging between 1% (dDG) to 12% (vCA1). Reactivation rates were not statistically different between the groups, and did not show any significant interactions between group, subregion, and/or axis (all *p*’s = 1.00) (Figure 2E).

### Susceptible animals show an increased number of vCA1 PV^+^ neurons that is recruited relatively less during remote fear memory recall

Given the influence of PV^+^ interneuronal activity on the excitability and firing behavior of surrounding neurons, we investigated PV^+^ activity during fear memory recall. For this we calculated PV^+^ ‘Activation Rate’ (AR) as the number of PV^+^cFos^+^ overlapping neurons divided by the number of PV^+^ neurons x 100%, reflecting the percentage of the total interneuronal PV^+^ population that was active during remote fear memory recall. In line with prior work indicating that Arc-expression in response to behavioral challenges is largely restricted to glutamatergic neurons (Vazdarjanova et al., 2006), the population of tdTomato-labeled (‘TRAPped’) neurons in this mouse line did not overlap with PV expression, which prevented us from also investigating relative activity of PV^+^ neurons during SEFL. Moreover, we also calculated the overall density of PV^+^ neurons to account for potential structural differences across groups.

PV^+^ ARs revealed a significant main effect of group (*F*(1,35.454) = 8.613, *p* = .006), together with a group x axis (*F*(1,35.681) = 9.045, *p* = .005), group x subregion (*F*(2,33.512) = 4.728, *p* = .016), and group x axis x subregion (*F*(2,33.512) = 5.172, *p* = .011) interaction effect (Figure 2F). Follow up tests revealed no significant group effects in the dorsal hippocampus (*p*’s > .543), but a significant main effect of group (*F*(1,18.823) = 14.541, *p* = .001) as well as a group x subregion interaction (*F*(2,15.804) = 7.056, *p* = .006) in the ventral hippocampus. This interaction effect was driven by significantly reduced PV^+^ activation in the vCA1 of susceptible mice (*p* = .001), but not other hippocampal subregions (both *p*’s > .340). The overall number of hippocampal PV^+^ neurons was not found to be significantly affected by group (*F*(1,35.978) = 1.533, *p* = .224). However, exploratory analyses to test whether the activity differences in vCA1 PV^+^ neurons were associated with different PV^+^ neuron numbers in susceptible mice, revealed a significantly higher PV^+^ neuron density in their vCA1 (*p* = .026), which was not observed in the other hippocampal subregions (all *p*’s > 0.227). These findings suggest that trauma susceptibility is associated with an increase in PV^+^ neurons in the vCA1, of which a relatively smaller part is active during remote fear memory recall.

### Susceptible mice display altered HDAC2 expression patterns in the ventral hippocampus

The intensity of HDAC2 fluorescence in engram and non-engram cells was measured to quantify HDAC2 expression within these neurons (Figure 3AB) (Toki et al., 2017, Ververis and Karagiannis, 2012). HDAC2 expression was dependent on hippocampal axis (*F*(1,207.453) = 159.497, *p* < .001), subregion (*F*(2,145.718) = 41.926, *p* < .001) and engram type (*F*(3,142.060) = 28.686, *p* < .001), but did not reveal a significant main effect of group (*F*(1,18.223) = 2.496, *p* = .131) (Figure 3C). Pair wise comparisons revealed that engram type effects were caused by significantly higher HDAC2 expression in memory encoding (tdTomato^+^; *p* < .001), recall (cFos^+^; *p* < .001) and reactivated (tdTomato^+^cFos^+^; *p* < .001) neurons, compared to non-engram cells, whereas the engram types amongst each other did not show overall differences in HDAC2 expression (all *p*’s > .320) (Figure 3CDE), suggesting histone acetylation is overall reduced in memory engram-related cells compared to non-engram cells.

**Figure 3.**
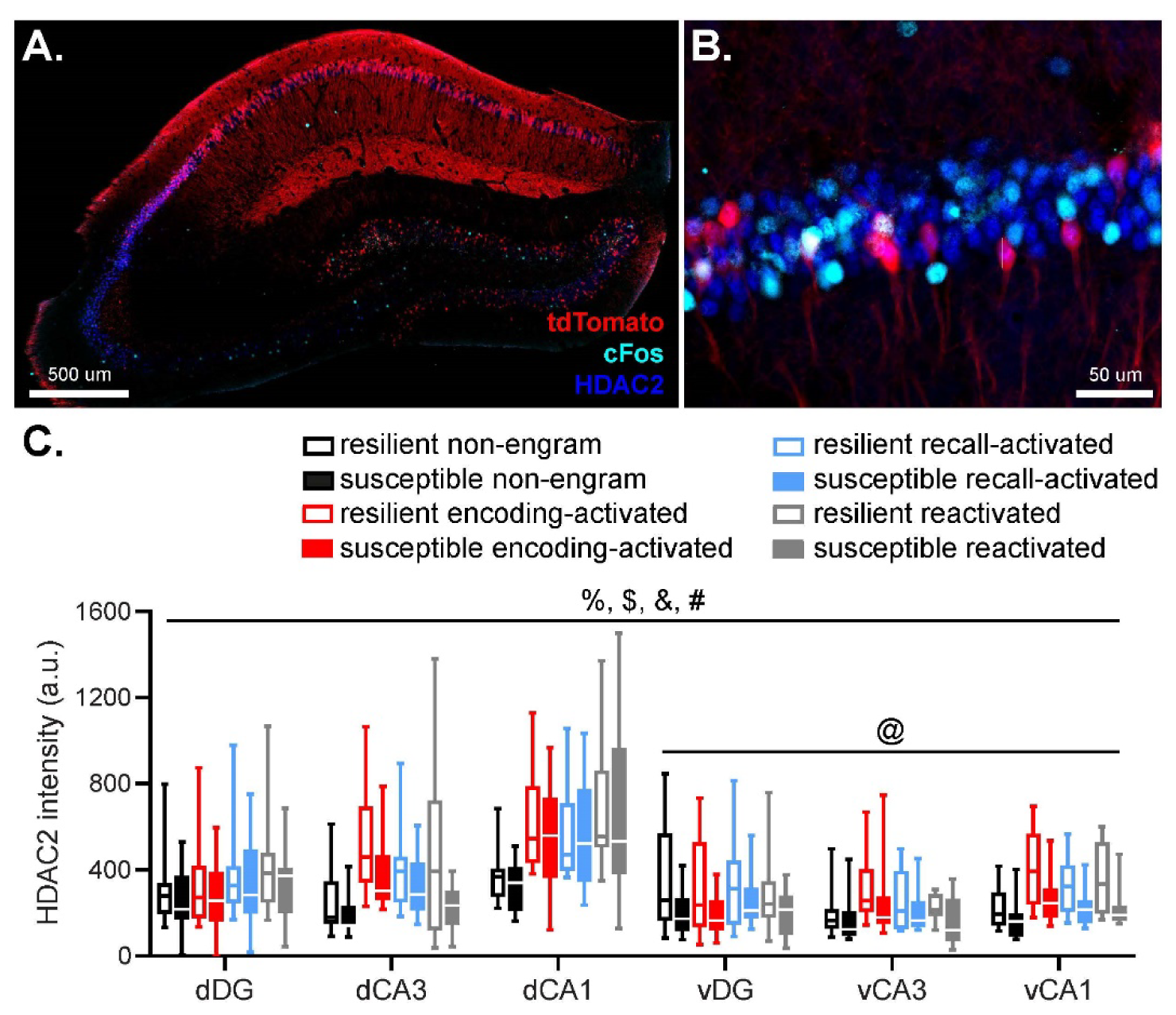
Hippocampal HDAC2 fluorescence in cells active during fear memory encoding (marked by tdTomato expression) and remote fear memory recall (marked by cFos expression) were assessed by immunohistochemistry (**AB**). Neurons involved in memory encoding (tdTomato^+^) and recall (cFos^+^), as well as reactivated (tdTomato^+^cFos^+^; indicated by arrows) neurons, were characterized by overall higher HDAC2 fluorescence than non-engram cells. HDAC2 levels in the ventral hippocampus were modulated by a subregion x group interaction, which seemed to be caused by a tendency towards lower HDAC2 levels in the vCA1 in susceptible animals (**CDE**). Data represent medians with interquartile ranges. %: p < .001, main effect of axis, $: p < .001, main effect of subregion, &: p < .001, main effect of engram type, #: p < .05, group x axis x subregion interaction, @: p < .05, group x subregion interaction.

Critically, we observed a significant group x hippocampal axis x subregion interaction in HDAC2 levels (*F*(2,145.718) = 3.467, *p* = .034). Follow up tests revealed no significant effects of group in the dorsal hippocampus (all *p*’s > .227), but a significant group x hippocampal subregion interaction (*F*(2,80.421) = 3.368, *p* = .039) in the ventral hippocampus. This interaction seemed to be caused by a tendency towards reduced HDAC2 levels in the vCA1 (*p* = .057) in susceptible compared to resilient mice, in the absence of differences in the vDG (*p* = .169) and vCA3 (*p* = .382).

### Susceptible animals show rather generic increases in hippocampal 5mC and 5hmC levels

The intensity of 5mC and 5hmC fluorescence in engram and non-engram cells was measured to determine the DNA methylation status of these neurons (Ramsawhook et al., 2017, Toki et al., 2017) (Figure 4AB). 5mC levels appeared to be modulated by hippocampal subregion (*F*(2,96.738) = 3.116, *p* = .049), engram type (*F*(3,46.479) = 27.426, *p* < .001) and group (*F*(1,14.271) = 5.324, *p* = .037), without a main effect of hippocampal axis (*p* = .567) (Figure 4C). Moreover, a significant group x engram type interaction was observed (*F*(3,46.479) = 3.389, *p* = .026), but no other significant interactions (all *p*’s >062). *Post hoc* comparisons revealed significant differences in 5mC levels between all types of engram cells, with memory encoding and reactivation cells displaying higher 5mC levels than non-engram cells (*p* < .001 and *p* = .004, respectively), whereas memory recall cells displayed significantly lower 5mC levels compared to non-engram cells (*p* = .005). Follow up tests on the group x engram type interaction revealed significant upregulation of 5mC levels of susceptible mice in memory encoding (*p* = .019), recall (*p* = .015) and non-engram cells (*p* = .029), without significant differences in reactivated cells (*p* = .107).

**Figure 4.**
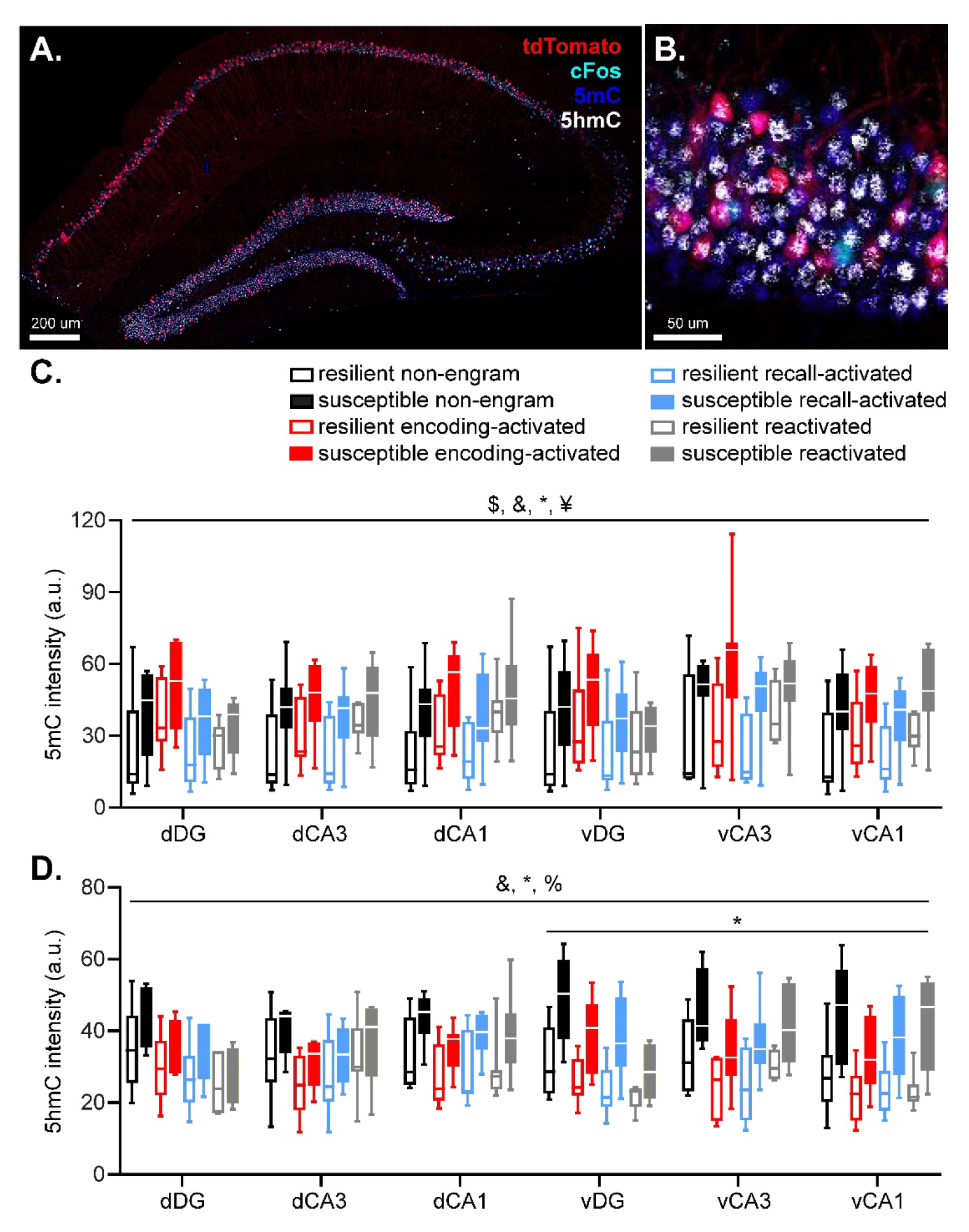
Hippocampal 5mC and 5hmC fluorescence in cells active during fear memory encoding (marked by tdTomato expression), remote fear memory recall (marked by cFos expression) and both were assessed by immunohistochemistry and compared to non-engram cells (tdTomato and cFos negative cells) (**AB**). 5mC levels were higher in encoding and recall cells compared to non-engram cells. In contrast, reactivated cells displayed lower 5mC levels than non-engram cells. Importantly, susceptible mice displayed higher 5mC levels in memory encoding, recall, and non-engram cells, compared to resilient mice, without significant differences in reactivated cells (**C**). 5hmC levels were lower in all types of engram cells compared to non-engram cells, and susceptible mice displayed higher 5hmC levels in the ventral hippocampus (**D**). Data represent medians with interquartile ranges. $: p < .05, main effect of subregion, &: p < .001, main effect of engram type, *: p < .05, main effect of group, ¥: p < .05, group x engram type, %: p < .001, group x axis interaction.

5hmC levels depended on engram type (*F*(3,97.708) = 24.770, *p* < .001) and group (*F*(1,16.006) = 6.837, *p* = .019), without a main effect of hippocampal axis (*p* = .457) or subregion (*p* = .431) (Figure 4D). Moreover, a significant group x axis interaction was observed (*F*(1,183.505) = 28.105, *p* < .001). All other interactions with group were non-significant (all *p*’s > .063). Pair wise comparisons revealed significantly lower 5hmC levels in all types of engram cells compared to non-engram cells (all *p*’s < .001), whereas the different type of engram cells (encoding, recall and reactivation) did not differ from each other (all *p*’s > .425). Follow up tests for the group x hippocampal axis interaction revealed that susceptible displayed significantly higher 5hmC levels in the ventral hippocampus (*F*(1,14.694) = 8.419, *p* = .011), but not dorsal hippocampus (*F*(1,14.190) = 3.295, *p* = .091).

While 5mC and 5hmC levels have been linked to decreased and increased gene expression respectively (Mendonca et al., 2014, Razin and Cedar, 1991), the 5hmC/5mC ratio might actually be most informative with regard to a cell’s gene expression profile, with high ratios coding increased gene expression (Mellen et al., 2012). Therefore, 5hmC/5mC ratios were calculated as well (Figure S2). 5hmC/5mC ratio data revealed a significant effect of engram type (*F*(3,71.552) = 65.954, *p* < .001), without any effects of hippocampal axis (*p* = .194), subregion (*p* = .540) or group (*p* = .210). Moreover, a significant group x engram type interaction was found (*F*(3,71.552) = 6.833, *p* < .001). Pair wise comparisons of 5hmC/5mC ratios revealed significantly lower ratio in engram vs. non-engram cells (encoding; *p* < .001, recall; *p* = .010, reactivation; *p* < .001), with encoding and reactivation cells displaying lowest ratio’s (both *p*’s < .001 compared to recall cells). This suggests that engram neurons are transcriptionally less active than neurons that are not incorporated into the engram, which is in line with previous studies marking increased DNA methylation in engram cells as a key mechanism in stabilizing memory engrams during memory consolidation (Gulmez Karaca et al., 2020). Follow up analyses on the group x engram type interaction however failed to indicate clear differences between susceptible and resilient mice (all *p*’s > .195).

## Discussion

Here, we tested the hypothesis that susceptibility to traumatic stress is characterized by interindividual differences in the trauma-related hippocampal memory engram and its epigenetic regulation. We examined potential alterations in the hippocampal memory engram for a stress-enhanced fear memory in mice that were susceptible and resilient to developing PTSD-like symptoms as a consequence of it. While no differences in the size of the neuronal ensemble activated during fear memory encoding were observed between the groups, susceptible mice displayed a smaller ensemble activated in the vCA1 upon remote fear memory recall, as well as higher PV^+^ neuronal density and a relatively lower activity of PV^+^ neurons in the vCA1 upon remote memory recall. Epigenetic data revealed rather generic than engram-specific differences across groups, with susceptible animals displaying lower hippocampal HDAC2 expression, as well as higher 5mC and 5hmC signal, without clear overall differences in 5hmC/5mC ratio.

Mice were classified as susceptible or resilient based on a compound score comprising multiple behavioral PTSD-like symptoms (i.e., impaired risk assessment, increased threat-induced anxiety, hypervigilance, impaired pre-pulse inhibition and higher activity during the inactive phase, potentially linking to sleep disturbances), rather than single behavioral features. Because of obvious limitations in capturing true PTSD-symptomatology in mice, these behavioral features should merely be considered as proxy’s for the complex behavioral symptomatology as observed in patients. Yet, the behavioral classification in which mice are categorized based on a compound score of symptomatology, resembles the situation in PTSD patients (Zoellner et al., 2014). No large differences were observed in how susceptible and resilient mice behaved during the encoding and recall of the fear memory. Whereas the susceptible mice showed a somewhat different temporal development of freezing during fear conditioning, no differences in freezing levels were observed during remote fear memory recall. Prior work has indicated that stress susceptible mice show exaggerated and extinction-resistant fear memory in a stress-enhanced cued fear learning paradigm (Sillivan et al., 2017). This fits observations of emotional hypermnesia in PTSD patients, as well as their strongly cue-based rather than context-specific recall of the trauma memory. Importantly, PTSD-like memory alterations also include contextual amnesia (Desmedt et al., 2015, Kaouane et al., 2012). We here implemented stress-enhanced *contextual* fear learning (Rau et al., 2005), making that one could speculate on impaired contextual fear memory recall canceling out excessive fear upon successful cue-induced recall in susceptible mice. Future studies need to confirm this speculation of a maladaptive fear memory in susceptible mice by dissociating both aspects of fear memory by re-exposing mice to partial fear reminders (cues vs context) only. Moreover, it would be valuable to include a regular (not stress-enhanced) fear learning group as control in future experiments, to both verify the effect of prior stress exposure overall and to determine whether these effects differ in resilient vs. susceptible mice.

Despite the absence of differential freezing behavior, we did find a significant reduction in vCA1 activity and a relative decrease in PV^+^ cell activation during remote fear memory recall in susceptible animals. Previous work has implicated the vCA1 in contextual fear memory (Kim and Cho, 2017, Maren and Fanselow, 1995, Rogers et al., 2006, Rudy et al., 2004, Zhu et al., 2014) and the subsequent contextual modulation of fear recall and expression (Orsini et al., 2011, Xu et al., 2016). Ventral CA1 neurons have been shown to convey contextual information through monosynaptic projections to the basolateral amygdala (Kim and Cho, 2017, 2020, Maren and Fanselow, 1995, Maren et al., 2013). As such, the reduction in the vCA1 ensemble activated during recall might reflect impaired functionality in the recall of contextual information, which may lie at the core of the context-nonspecific recall of trauma memories as observed in PTSD (Asok et al., 2018, Liberzon and Abelson, 2016, Zinn et al., 2020) as well as the reported contextual amnesia (Desmedt, 2021). As we did not find any differences in engram size during fear memory encoding (and reactivation), data suggest that initial memory encoding is not different between groups, but it is rather the (systems) consolidation process during which differences arise. The memory engram is not static, but rather dynamic over time, reorganizing both within and across brain regions (Davis and Reijmers, 2018, M. E. Wang et al., 2015, Ziv et al., 2013), ultimately resulting in different storage sites of the memory following its consolidation (Davis and Reijmers, 2018, Frankland et al., 2004, Tayler et al., 2013, Tonegawa et al., 2015, Wang et al., 2015, Ziv et al., 2013). This is especially relevant as we employed a remote recall paradigm, whereas most previous studies focused on more recent memory recall and might explain why we only observed differential vCA1 activity and not reactivity during recall.

Parvalbuminergic network plasticity has been shown critical in the regulation of learning (Donato et al., 2013), with PV^+^ interneurons contributing to memory consolidation by stabilizing functional connectivity patterns among CA1 neurons (Ognjanovski et al., 2017) and mediating coherent hippocampal-neocortical communication (Xia et al., 2017). We observed a higher number of PV^+^ neurons in the vCA1 of susceptible animals as well as a smaller portion of these being activated during memory recall. Seemingly contradictory to our findings, prior research has reported on a loss of PV^+^ neurons following chronic stress (Czeh et al., 2015, Holm et al., 2011), although this is not consistently reported (Holm et al., 2011, Nowak et al., 2010). Yet, notably, previous studies have ignored interindividual differences in terms of anxiety-like behavior, posing the possibility that this observed reduction in PV^+^ density may actually reflect an adaptive response to stress. Alternatively, the differences between susceptible and resilient mice may have been present already before PTSD induction, and as such do not reflect a differential effect of trauma itself. The lower recruitment of these neurons in susceptible mice may indicate a compensatory effect, resulting in similar absolute activity levels of the total PV^+^ population in both susceptible and resilient animals. Regardless, these alterations in PV^+^ interneuron presence and recruitment might relate to disrupted consolidation of the traumatic memory in PTSD (Parsons and Ressler, 2013), proposing it as a target for dedicated future studies.

Hippocampal HDAC2 expression was higher in engram compared to non-engram cells and reduced in susceptible compared to resilient animals. Histone acetylation is most robustly associated with promoting memory formation. It is increased following neuronal activity, and promotes a chromatin structure permissive to gene transcription (Eberharter and Becker, 2002), necessary for synaptic plasticity (Marco, 2022). HDACs, in particular HDAC2, induce the removal of acetyl groups, suppressing gene transcription, and their pharmacological or genetic inhibition was found to facilitate learning and memory (Guan et al., 2009) and improve extinction learning (Vecsey et al. 2007, Morris et al., 2013, Graff et al., 2014). Our finding of increased HDAC2 levels in engram vs. non-engram cells seems to be at odds with these reports. Yet, one could speculate that plasticity should be suppressed once a memory is formed, with memory-related gene silencing serving to stabilize the memory engram (Alberini and Kandel, 2014). This interpretation is supported by our findings in terms of DNA methylation patterns, with engram cells having overall higher levels of 5mC (generally suppressing gene transcription) and lower levels of 5hmC (typically increasing gene transcription), decreasing the 5hmC/5mC ratio, suggesting an overall decrease in transcriptional activity within engram neurons. Prior reports implicating DNA methylation in stabilizing engrams during consolidation and aiding successful memory recall support this notion (Gulmez Karaca et al., 2020). As such, reduced HDAC2 levels as observed in susceptible mice may indicate a less stable fear memory engram. Whereas this interpretation might fit with the readily re-activated trauma memory in PTSD, it is at odds with behavioral observations of a trauma memory in patients that is very rigid, and less sensitive to extinction. However, our findings are in line with prior reports on HDAC2 downregulation following acute stress being related to increased stress susceptibility (Karnib et al., 2019) and a stronger fear memory (Takei et al., 2011). In terms of methylation, we found susceptible animals to be characterized by rather generic increases in hippocampal 5mC and 5hmC levels, both in engram and non-engram cells. As these markers are inversely related to gene expression and their ratio was not consistently affected, we conclude that both groups, despite the slight differences in hippocampal methylation profile, do likely not differ in terms of overall gene expression as a consequence of this. Prior research has reported on changes into hippocampal global methylation levels as a consequence of stress exposure, with both increases (Kashimoto et al., 2016) and decreases (Rodrigues et al., 2015; Sales et al., 2020) being reported. We add to this existing literature by relating methylation patterns to interindividual differences in stress susceptibility, which match reports on increased global methylation in PTSD patients (Smith et al., 2011). Yet, future studies should investigate these differences in more detail, by assessing specific methylation sites, as well as potential mediators of these differences (e.g., DNA methyltransferases). Moreover, as the observed epigenetic differences cannot readily explain the reduced number of cFos expressing cells upon memory recall in susceptible mice, it would be worthwhile investigating other epigenetic regulators. One of these might be HDAC5, which was previously found to be upregulated post-trauma in the bed nucleus stria terminalis of susceptible animals in this same mouse model (Lebow et al., 2012).

Some limitations should be noted. Firstly, assessment of the memory engram related to memory encoding was restricted to glutamatergic neurons in the ArcTRAP mice (Vazdarjanova et al., 2006), leaving the role of GABAergic neurons in the engram and traumatic stress susceptibility to be elucidated. We preferred ArcTRAP mice over the available FosTRAP mice based on superior labeling sensitivity in the hippocampal CA3 and CA1, which are typically devoid of labeled cells in the FosTRAP mouse lines (Dirven et al., 2022b, Guenthner et al., 2013). Secondly, the ArcTRAP line has substantial background labeling (i.e., fluorescent tagging of neurons in the absence of tamoxifen) in the hippocampal DG (Guenthner et al., 2013), which may explain why we did not recapitulate prior findings of peri-trauma DG activation being predictive of fear memory generalization and stress susceptibility in general (Dirven et al., 2022b, Lesuis et al., 2021). Furthermore, tdTomato-labeling captured both neuronal activity during the initial stressor and subsequent fear learning experience. Future studies should separate these two episodes by making use of 4-hydroxytamoxifen injection, restricting the labeling period. Moreover, while we assume that the tdTomato-tagged and cFos-labelled neurons represent the fear memory, it will require experimental manipulation of these populations to show that their activity is necessary and/or sufficient for memory expression. Finally, while immunofluorescence of epigenetic markers is more often used to draw preliminary conclusions about changes in transcriptional processes (Chouliaras et al., 2013, Demyanenko and Uzdensky, 2019), it is not possible to draw a one-to-one relationship between the observed differences in HDAC2, 5mC and 5hmC levels and actual alterations in histone acetylation and gene expression. Different studies have shown transcriptional alterations in response to stress (Floriou-Servou et al., 2018), and in PTSD specifically (Girgenti et al., 2021, Zhang et al., 2021), but it would require future studies to causally link such changes to the alterations in histone acetylation, DNA methylation and hydroxymethylation that have been observed.

Concluding, we have shown PTSD-like symptomatology in mice to be related to alterations in remote fear recall-induced activation and PV^+^ interneuronal activity - as well as overall PV^+^ density - in the ventral CA1. These findings propose an important role for aberrant remote fear memory recall, resulting from an altered (systems) consolidation process, in mediating traumatic stress susceptibility. Future assessments should investigate whether this can also translate into an aberrant behavioral manifestation of the fear memory. Epigenetically, we found marked differences in HDAC2 expression and DNA methylation and hydroxymethylation between susceptible vs. resilient mice, suggestive of net higher hippocampal transcriptional activity. These changes were however not restricted to neurons involved in the memory engram, indicating epigenetic changes throughout the entire hippocampus as an important target for further research into the pathophysiology of PTSD. These overall alterations could potentially contribute to deviations in memory consolidation by destabilizing hippocampal memory representations, although future research is needed to determine such causal relationship.

## Acknowledgements

M.J.A.G.H. was supported by Veni Grant 863.15.008 and Vidi Grant 203.028, and J.R.H. by Vidi Grant 864.10.003 awarded by the Netherlands Organization for Scientific Research.

## Data Availability Statement

All experimental data and analyses tools will be shared upon request.

