## Supplementary Figures for "Hippocampal trauma memory processing conveying susceptibility to traumatic stress"

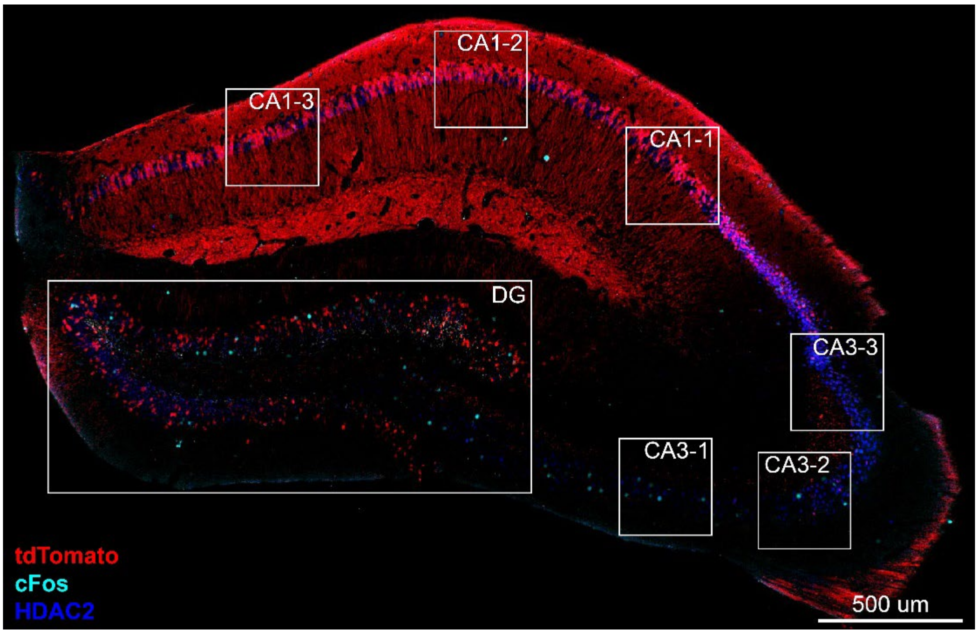


**Figure S1.** Hippocampal HDAC2 fluorescence was assessed in 40x microscopic frames. For the DG, multiple frames were stitched to obtain a photo of the entire structure. For the CA3 and CA1, three representative photos each were taken in locations consistent across slices and animals.

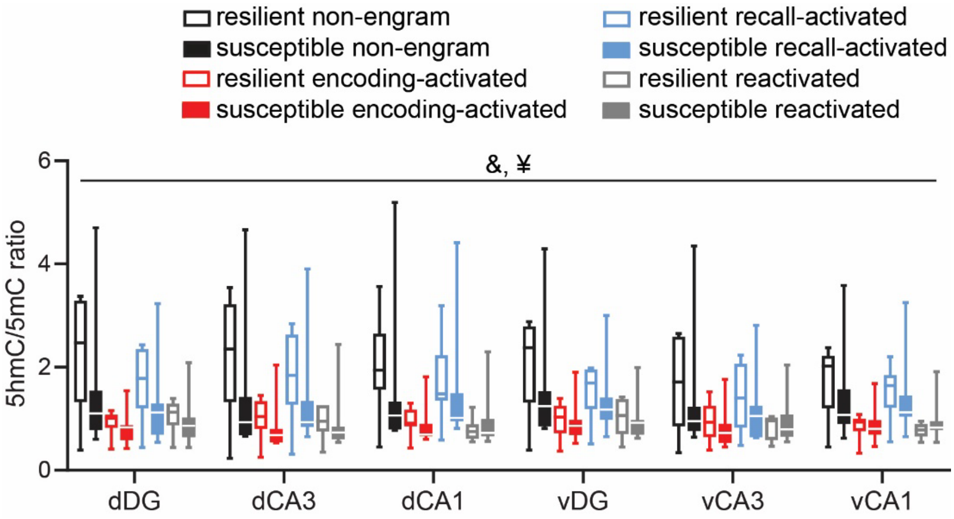


**Figure S2.** Hippocampal 5hmC/5mC fluorescence ratio in cells active during fear memory encoding (marked by tdTomato expression) and remote fear memory recall (marked by cFos expression) as assessed by immunohistochemistry. All types of engram neurons displayed lower ratio than non-engram cells, with encoding-activated (tdTomato$^+$) and reactivated (tdTomato$^+$cFos$^+$) neurons displaying lowest 5hmC/5mC ratios, indicative of a relative decrease in gene transcription. Data represent medians and interquartile ranges. &: $p$ < .001, main effect of engram type, ¥: $p$ < .001, group x engram type interaction.
